# Shh signaling directs dorsal ventral patterning in the regenerating *X. tropicalis* spinal cord

**DOI:** 10.1101/2024.10.18.619160

**Authors:** Avery Angell Swearer, Samuel Perkowski, Andrea Wills

## Abstract

Tissue development and regeneration rely on the deployment of embryonic signals to drive progenitor activity and thus generate complex cell diversity and organization. One such signal is Sonic Hedgehog (Shh), which establishes the dorsal-ventral (D/V) axis of the spinal cord during embryogenesis. However, the existence of this D/V axis and its dependence on Shh signaling during regeneration varies by species. Here we investigate the function of Shh signaling in patterning the D/V axis during spinal cord regeneration in *Xenopus tropicalis* tadpoles. We find that neural progenitor markers Msx1/2, Nkx6.1, and Nkx2.2 are confined to dorsal, intermediate and ventral spatial domains, respectively, in both the uninjured and regenerating spinal cord. These domains are altered by perturbation of Shh signaling. Additionally, we find that these D/V domains are more sensitive to Shh perturbation during regeneration than uninjured tissue. The renewed sensitivity of these neural progenitor cells to Shh signals represents a regeneration specific response and raises questions about how responsiveness to developmental patterning cues is regulated in mature and regenerating tissues.

## Introduction

Regeneration is often understood through the lens of development, as both processes must generate and pattern complex tissue. During embryogenesis, positional identity in the vertebrate spinal cord is established through opposing gradients of secreted dorsal bone morphogenetic protein (BMP) and ventral Sonic hedgehog (Shh) signals, which are interpreted through a transcription factor code to yield distinct dorsal, intermediate, and ventral neural progenitor cell (NPC) domains^1,2^. More specifically, this gradient of Shh is secreted from the ventrally-located notochord and floor plate and is canonically interpreted through Gli transcription factors. In the presence of the Shh ligand, these factors are converted from transcriptional repressors to activators, and drive expression of ventral NPC identities and repression of dorsal identities^3^. Each unique NPC domain along the Dorsal/Ventral (D/V) axis later gives rise to a similarly spatially restricted domain of post-mitotic neurons, which work together to mediate the many types of afferent and efferent neural impulses of the spinal cord^4,5^. The basic features of this developmental patterning mechanism are fundamentally conserved in vertebrates, with fish, frogs, chicks, mice, and others all showing disruptions of spinal cord patterning when BMP or Shh are perturbed^6–13^.Shh has also been shown to act in a mitogenic role to support NPC proliferation in both the embryonic and mature CNS in the chick and mouse^14–16^.

In animals that can regenerate, one might intuitively expect that regeneration of spinal cord cell types would re-use the same developmental mechanisms to pattern lost tissue. However, there are examples that both support and oppose this paradigm. Across the vertebrate phylum, BMP and Shh are understood to be required for regeneration, as the absence of these signals leads to incomplete regeneration in zebrafish, *Xenopus*, axolotls, newts and lizards^17–24^. The role of these signals, specifically Shh, in conferring positional identity to the regenerated neural progenitors varies among these species. In axolotls (*Ambystoma mexicanum*), Shh specifies ventral neural stem/progenitor cell domains during regeneration, a process that can be disrupted by inhibition of Shh signaling^21^. However, axolotls also appear to take advantage of the existing pattern in the mature spinal cord, as most regenerated daughter cells occupy the dorsoventral position of their parent^25^. Thus, both the *de novo* establishment of positional identity by Shh and the positional identity of pre-existing progenitors may contribute to D/V patterning in the regenerated spinal cord. By contrast, in green anole lizards (*Anolis carolinensis*), the spinal cord regenerates as a ubiquitously Shh+ ependymal tube, in which Shh serves to specify surrounding mesoderm but has no proven influence in generating a D/V axis^24^. Thus the role of Shh in directing D/V patterning in the regenerated spinal cord is less universal than its role in the developing spinal cord.

*Xenopus* tadpoles occupy a unique niche among regeneration-competent animals because they are capable of functional spinal cord regeneration prior to metamorphosis, but lose that ability as adults^26^. In *X. laevis*, Shh is expressed in the regenerating notochord following injury, and this signal is necessary for the full regeneration of not only the spinal cord, but also the muscle and notochord^20^. However, it is not clear that the requirement for Shh in regeneration derives from its canonical functions through Gli transcription factors. Most notably, knockdown of Gli2 and treatment with the Gli1/2 inhibitor GANT61 does not impair global regeneration in *X. laevis*. Rather, a non-canonical Shh pathway of Ca^2+^-dependent activation and coordination of proliferation in neural and muscle progenitor cells is thought to direct the Shh-dependent aspects of regenerative growth in the tail^27^. This reported lack of function for Gli1/2 during regeneration could be due to the absence of D/V patterning during regeneration, leading us to question whether these D/V domains are regenerated and if Shh retains any developmental patterning function during regeneration in *Xenopus*.

Here we interrogated the basis for D/V patterning during regeneration of the spinal cord in *Xenopus tropicalis*, a diploid relative of *X. laevis* that shares many of the latter’s regenerative features and mechanisms^28^. We specifically sought to determine whether the regenerating spinal cord recapitulates the D/V organization of the embryonic spinal cord by examining expression of dorsal, intermediate, and ventral NPC markers. Next, we asked whether this organization is dependent on Shh through gain and loss of function perturbations. Finally, we sought to identify whether positional identity in the uninjured spinal cord is plastic or fixed, by asking whether perturbations of Shh change the balance of positional identity in uninjured tadpoles as compared to regenerating tadpoles.

## Results

### Dorsal, intermediate, and ventral NPC domains exist in the uninjured, regenerative-stage spinal cord

Our first goal was to confirm that NPCs were organized into spatially restricted dorsal, intermediate, and ventral domains in uninjured, regenerative-stage *X. tropicalis* tadpoles. To this end, we identified an antibody against two transcription factor paralogs, Msx1 and Msx2, which have conserved expression in dorsal NPCs of the developing spinal cord downstream of BMP signaling^29–31^. We then selected two transcription factors downstream of Shh signaling, Nkx6.1 and Nkx2.2, as intermediate and ventral markers, respectively. These genes are known to be expressed in intermediate and ventral domains of the developing spinal cord in *Xenopus*^12,32,33^. We performed immunofluorescence for Msx1/2, Nkx2.2, and Nkx6.1 to assess expression of these markers in the uninjured spinal cord of regenerative Stage 46 uninjured tadpoles (Figure 1). In lateral views and transverse sections, each marker was found to be expressed in their expected D/V domains.

**Figure 1.**
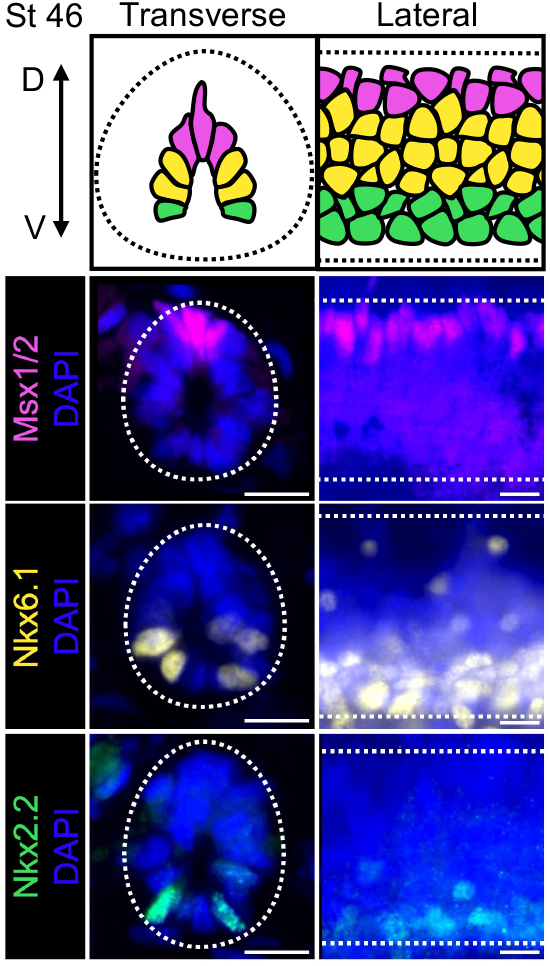
Dorsal, intermediate, and ventral NPC domains can be visualized in the uninjured, regenerative-stage spinal cord. Top, diagram of Stage 46 uninjured regenerative-stage spinal cord. Bottom, transverse and lateral images visualizing dorsal markers Msx1 and Msx2 (Msx1/2: 4G1), intermediate marker Nkx6.1 (F44A10), and ventral marker Nkx2.2 (74-5A5). D = Dorsal, V = Ventral, dotted line indicates spinal cord boundary. Scale bars = 10 µm.

### Dorsal, intermediate, and ventral NPC domains are present within the regenerating *X. tropicalis* spinal cord

Having confirmed D/V NPC domains in uninjured tissue, we next asked if this organization is recapitulated in the regenerating spinal cord. It was previously reported that some other regenerative animals such as the axolotl do regenerate regionalized D/V domains, but others, like the green anole lizard, only regenerate a ventralized ependymal tube. To determine if these D/V NPC domains are present during *Xenopus* regeneration, we amputated the posterior third of the tail of stage 41 tadpoles, as we have previously^34^. Tadpoles were allowed to regenerate for 2, 3, or 4 days post amputation (dpa), before they were collected and stained for Msx1/2, Nkx6.1, or Nkx2.2 in transverse section and whole mount preparations (Figure 2A). We found that throughout the surveyed time period, Msx1/2, Nkx6.1, and Nkx2.2 are regionally localized into dorsal, intermediate and ventral domains in the regenerating spinal cord (Figure 2B-D). In order to quantify this NPC localization pattern, we measured the position of cells expressing each marker in whole mount images and normalized this position to spinal cord width or length (Figure 2E). Localization of Msx1/2+ cells are clearly biased to the dorsal spinal cord, with Nkx6.1+ cells positioned intermediately and Nkx2.2+ cells ventrally. Interestingly, we identified not only a D/V NPC axis, but also A/P regionalization of these markers during regeneration. Specifically, Msx1/2+ cells were more strongly regionalized posteriorly, while Nkx2.2+ cells were more strongly regionalized anteriorly (Figure 2E). From these data, our principal conclusion was that the regenerated spinal cord is indeed patterned along the D/V axis, eventually recovering a similar organization to its pre-injury condition. In this regard, the *X. tropicalis* spinal cord more closely resembles that of the axolotl, in which NPCs are also organized dorsal-ventrally, rather than the anole, which is radially ventralized^24^.

**Figure 2.**
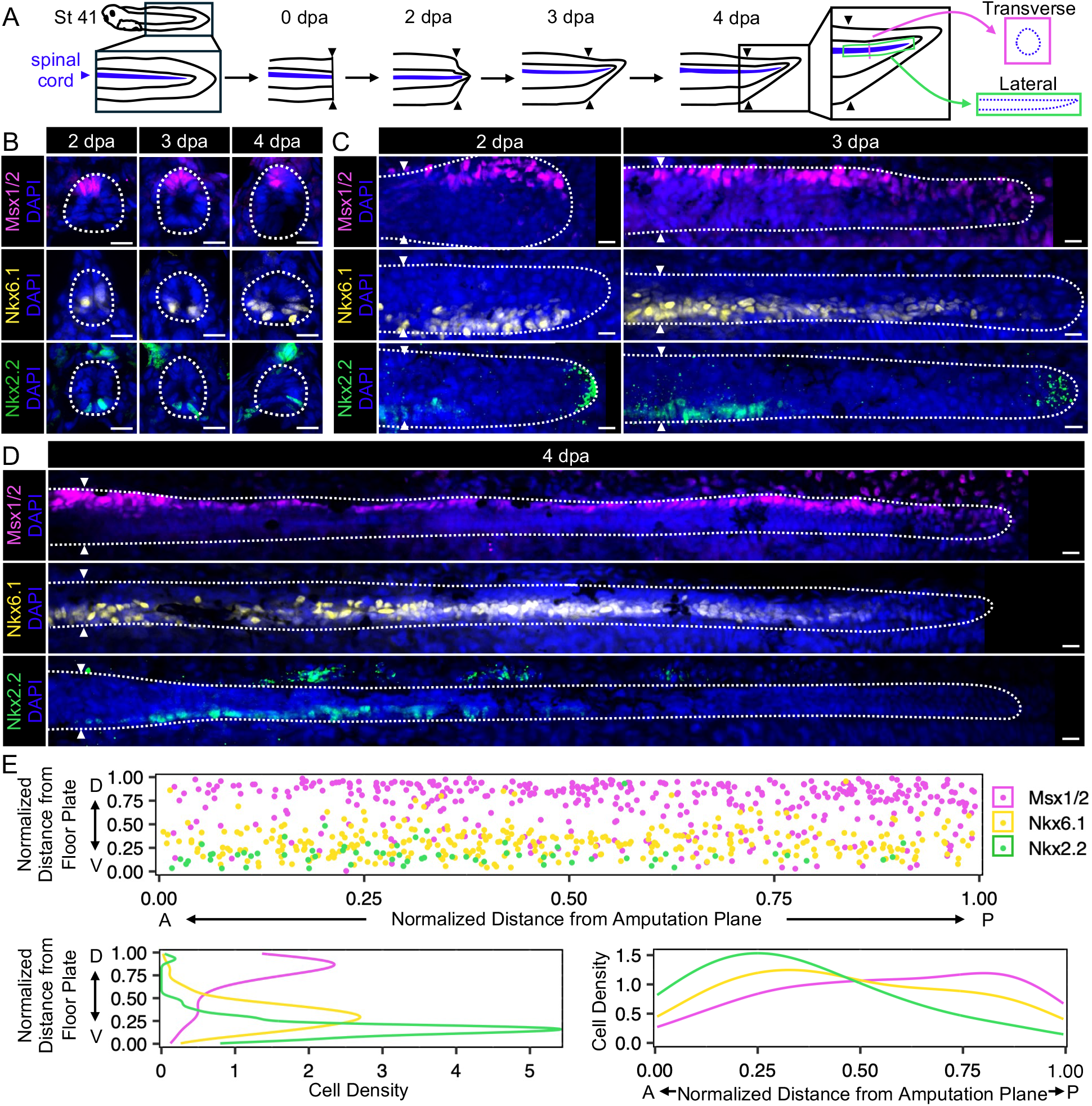
Dorsal, intermediate, and ventral NPC domains are present within the regenerating X. tropicalis spinal cord. **A**. Experimental design for sample collection post amputation. **B**. Images of transverse sections visualizing dorsal marker Msx1/2, intermediate marker Nkx6.1 and ventral marker Nkx2.2 at 2, 3, or 4 dpa. **C-D**. Images of whole mount tadpoles visualizing Msx1/2, Nkx6.1 and Nkx2.2 at 2, 3, or 4 dpa. **E**. Top, scatter plot showing cumulative cells based on their distance from floor plate or amputation planes, normalized to spinal cord width or length. Bottom, smoothed cell density plots illustrating regionalized distribution of D/V Markers over the D/V axis and A/P axis, respectively. For all images in this figure, paired triangles represent amputation plane and dotted lines represent spinal cord boundary. n = 17-20 per Marker. D = Dorsal, V = Ventral. Scale bar = 10 µm.

### Shh signaling is necessary and sufficient for D/V NPC patterning during regeneration

We next asked what mechanisms help specify NPC positional identity along the D/V axis during the rapid progenitor proliferation and cell movements that accompany regeneration. Here we hoped to discriminate between two possible mechanisms for D/V patterning during regeneration. By one mechanism, D/V domains could be maintained via the simple mechanism of clonal restriction to parental domains. In this model, each newly born NPC would retain the positional identity of its parent, and morphogens such as BMP or Shh would be dispensable for positional identity in the regenerate. Alternatively, regenerative neural progenitors could be multipotent with respect to positional identity, and extrinsic signals such as Shh would be required to reestablish D/V progenitor domains. Research in axolotls has shown that both mechanisms can be active during regeneration^21,25^. In *Xenopus*, both Shh and BMP are required for regeneration^20,23^. However, that does not necessarily imply that their D/V patterning functions are specifically required, since both these signals have pleiotropic effects that also include regulation of cell proliferation. In light of the non-canonical functions for Shh during regeneration, the role for Shh in D/V patterning was particularly called into question^27^. Therefore, we set out to directly ask whether perturbation of Shh disrupted D/V patterning in the regenerate.

Before testing whether Shh is essential for the patterning of NPCs during regeneration in *X. tropicalis*, we first asked where and when *shh* is expressed during regeneration. Visualization of *shh* transcripts via *in situ* hybridization showed continued low expression of *shh* in the st. 41 uninjured tadpole tail. Expression of *shh* is found within the regenerating notochord starting as early as 8 hpa, and increases in intensity in the regenerating notochord and spinal cord floor plate starting at 2 dpa (Figure 3A). This early expression of *shh* within the regenerating notochord matches with previously reported results in *Xenopus laevis* and zebrafish^17,20^. The expression of *shh* along with floor plate marker *foxa2* (Figure S1) in the spinal cord is notable, because previous studies done in *Xenopus laevis only* observed *shh* expression in the notochord, and not the floor plate, during regeneration^20^. This suggests a possible divergence between the two species. Overall, these results place *shh* expression at the right time and place to have an early effect on the D/V axis, as expected from previous studies in *Xenopus laevis* and other vertebrates.

**Figure 3.**
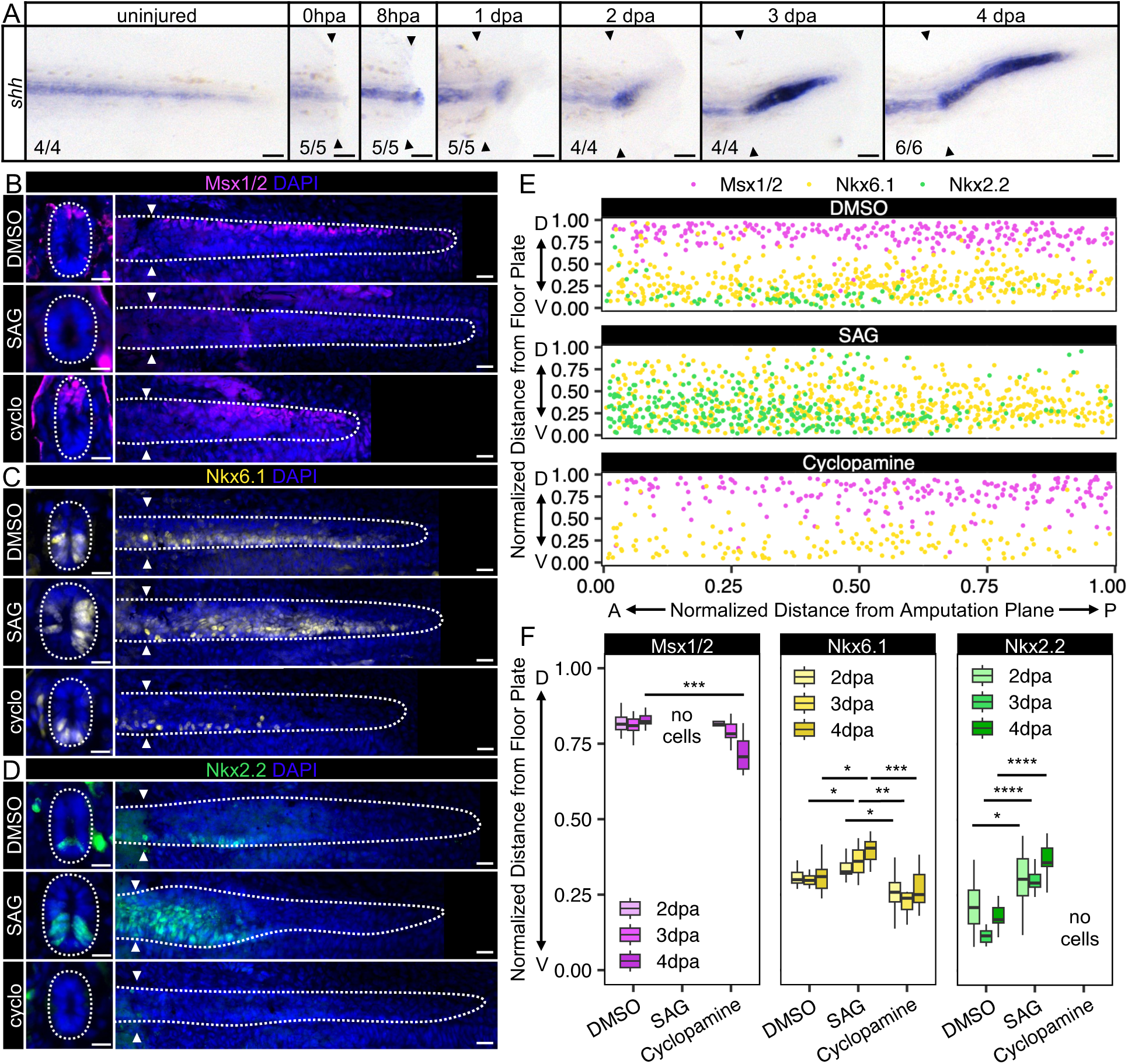
Shh signaling is necessary and sufficient for D/V NPC patterning during regeneration. **A**. Images of *in situ* hybridization visualizing *shh* expression at 0 hours post amputation (hpa), 8 hpa, 1 dpa, 2 dpa, 3 dpa, and 4 dpa. n/n is noted. Scale bar = 100 µm. **B-D**. Images of transverse sections (left) or lateral views (right) of tadpoles treated with DMSO, Shh agonist SAG or Shh inhibitor cyclopamine (“cyclo”) for 3 dpa and stained for Msx1/2 (**B**), Nkx6.1 (**C**), Nkx2.2 (**D**). Scale bar = 10 µm. n = 10-17 tadpoles per condition. **E**. Scatter plot showing cumulative cells based on their distance from floor plate or amputation planes, normalized to spinal cord width or length after DMSO, SAG, or cyclopamine treatment for 3 dpa. D = Dorsal, V = Ventral. n = 10-17 tadpoles per condition **F**. Quantification of average normalized distance from floor plate after treatment with DMSO, SAG, or cyclopamine treatment over 2, 3, and 4 dpa. n = 6-17 tadpoles per condition. Significance was determined by one-way ANOVA followed by Tukey’s multiple comparisons test or by pairwise two-sample Wilcoxon tests ( ^*^ p<0.05, ^**^ p<0.01, ^***^ p<0.001, ^****^ p<0.0001). For all images in this figure, dotted white lines represent spinal cord boundary and paired triangles represent amputation plane.

To test Shh’s functional role in shaping the D/V axis during regeneration, we used the hedgehog pathway inhibitor cyclopamine and agonist SAG to modulate Shh activity. Cyclopamine and SAG have both been used to inhibit Shh signal transduction in *Xenopus* and other vertebrates^27,33,36–38^. Because high doses of cyclopamine completely prevent spinal cord and tail regeneration, we performed a dose curve to identify doses that did not globally impair regeneration of the spinal cord. This allowed us to isolate the effect of Shh modulation on spinal cord patterning (Figure S2).

Following treatment with cyclopamine, the spinal cord exhibited an expansion of dorsal Msx1/2+ cells, reduction in the number of intermediate Nkx6.1+ cells, and complete absence of ventral Nkx2.2+ expression in NPCs. Treatment with cyclopamine did not significantly affect the average position of Nkx6.1+ cells in lateral view, but after refreshing the media every 24 hours we did observe a ventral shift in expression (Figure 4B). The opposite result was seen following treatment with SAG: Nkx2.2+ and Nkx6.1+ domains expanded significantly, while Msx1/2 expression was lost (Figure 3B-F). These two complementary results demonstrate that Shh is required for normal repatterning of NPCs after injury, outside of its reported non-canonical mitogenic function in supporting stem cell proliferation.

**Figure 4.**
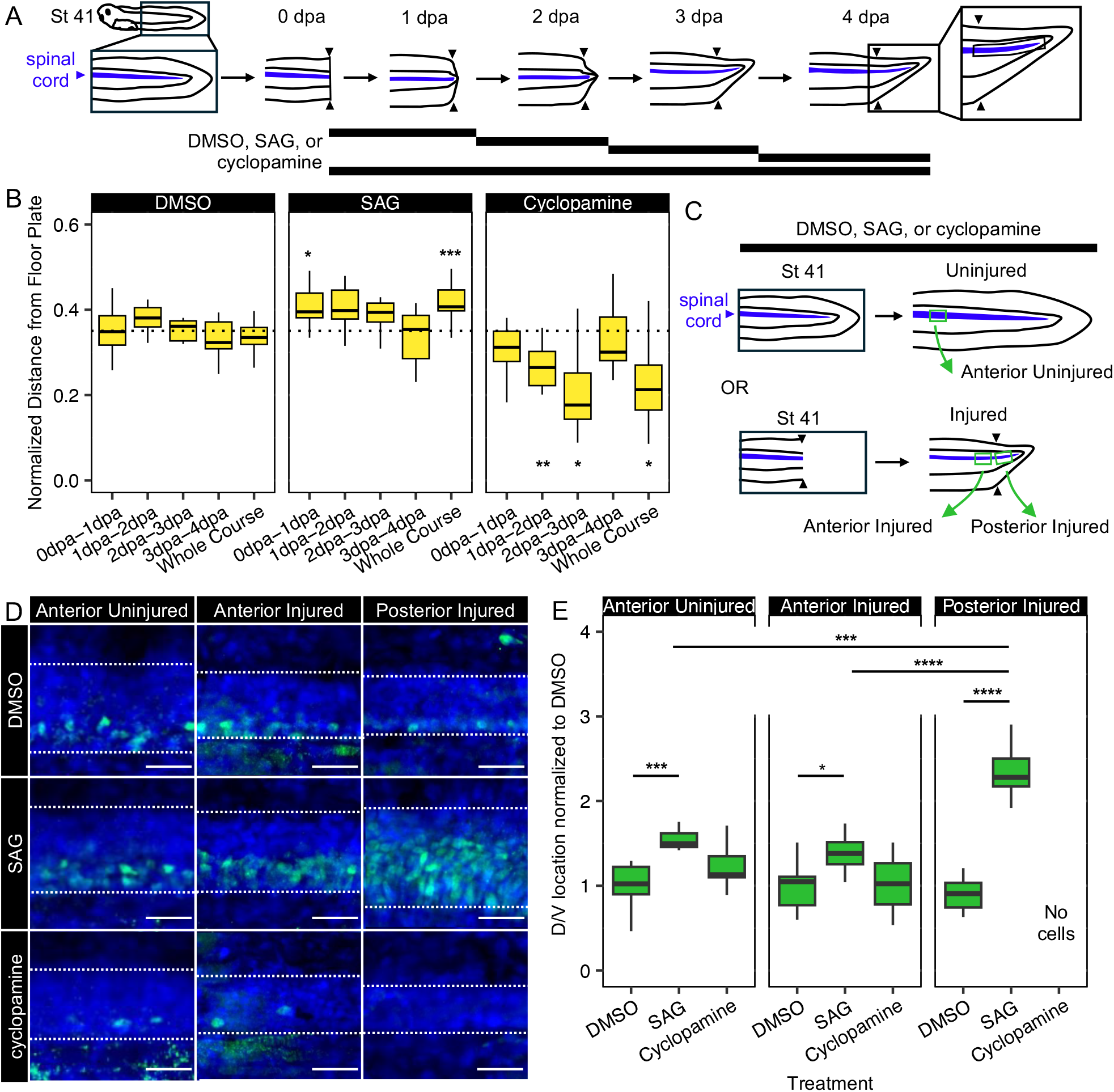
Regenerating NPCs are more sensitive to Shh modulation before 3 dpa and compared to uninjured cells. **A**. Experimental design for DMSO, SAG, or cyclopamine treatment during regenerative timepoints. **B**. Quantification of average normalized distance of Nkx6.1+ cells from the floor plate after treatment with DMSO, SAG, or cyclopamine treatment during specific regenerative timepoints, where the dotted black line represents average DMSO normalized D/V location across all timepoints. n = 9-15 tadpoles. Significance was determined by one-way ANOVA followed by Tukey’s multiple comparisons test or by pairwise two-sample Wilcoxon tests ( ^*^ p<0.05, ^**^ p<0.01, ^***^ p<0.001, ^****^ p<0.0001), against to comparative DMSO timepoint. **C**. Experimental design for collection of Anterior Uninjured, Anterior Injured, and Posterior Injured tissue after DMSO, SAG, or Cyclopamine treatment for 3 dpa. **D**. Images showing lateral views visualizing ventral marker Nkx2.2 within the Anterior Uninjured, Anterior Injured, or Posterior Injured spinal cord after treatment with DMSO, SAG, or cyclopamine. Dashed white lines represent spinal cord boundary. Scale bar = 10 µm. **E**. Quantification of (**D**), with D/V location normalized to DMSO. n = 8-11 tadpoles per condition. Significance was determined by pairwise two-sample Wilcoxon tests ( ^*^ p<0.05, ^**^ p<0.01, ^***^ p<0.001, ^****^ p<0.0001).

**Figure 5.**
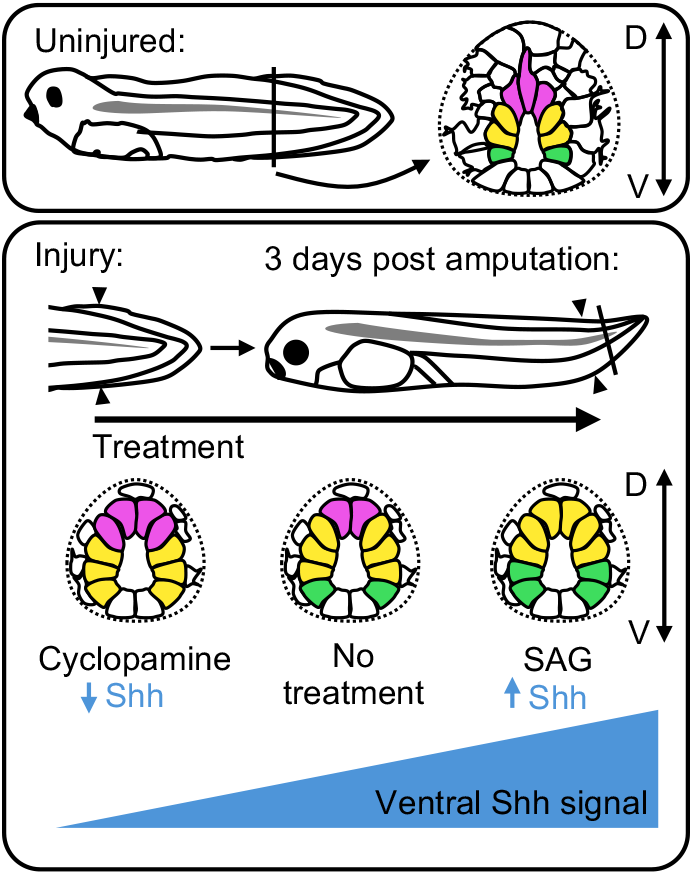
Shh drives NPC patterning after spinal cord amputation. In uninjured tadpoles, neural progenitor cells (NPCs) are dorsal/ventrally (D/V) patterned. By 3 days post amputation, this patterning regenerates, driven by ventral Shh signaling in a recapitulation of development. Inhibition of Shh using cyclopamine dorsalizes the regenerating spinal cord, whereas increasing Shh signaling with SAG ventralizes the spinal cord.

### Restricted window for sensitivity to Shh perturbation post amputation

To determine how the Shh signal response may change over the course of regeneration, we performed 1 day-restricted treatments of cyclopamine and SAG from 1 to 4 days post amputation and stained for the intermediate marker Nkx6.1 at 4 dpa. We chose Nkx6.1 because of its broad range of expression in lateral views (Figure 4A). We observed that treatment with SAG during the initial 1 day period after injury resulted in a significant expansion of the Nkx6.1+ domain. SAG treatment after 1 dpa resulted in expansion of the Nkx6.1 domain, but no significant change in average dorsal-ventral position of cells. Treatment with cyclopamine only resulted in a significant change in the Nkx6.1+ domain when tadpoles were treated during 1-2 and 2-3 dpa. We attribute the observed lack of sensitivity to cyclopamine treatment within the first day after injury to the return of normal Shh signal transduction levels after removal of cyclopamine from the media. Treatment with either drug had no effect on the position of the Nkx6.1+ domain after 3 days post amputation (Figure 4B, S3). From these results we find that the Nkx6.1 expression domain is sensitive to Shh perturbation immediately after injury, but loses sensitivity after three days of regeneration.

### Regenerating cells have altered response to Shh perturbation compared to uninjured cells

In the absence of injury, *shh* is still weakly expressed in the notochord and floor plate. This made us question how much Shh signaling might be contributing to the ongoing maintenance of NPC identity in the uninjured tadpole. To address this question, we asked whether regionalized NPCs changed their domain boundaries when Shh was perturbed in uninjured tadpoles, and compared this effect with Shh perturbation in anterior spinal cord tissue of injured tadpoles (tissue anterior to the amputation plane) and with Shh perturbation in regenerating spinal cord tissue (tissue posterior to the amputation plane). Using the same protocol used for the injured tadpoles, we treated uninjured stage 41 tadpoles with SAG or cyclopamine for three days. We then stained for ventral marker Nkx2.2, chosen because of its high sensitivity to Shh perturbation (Figure 4C). We found that the Nkx2.2+ domain remains sensitive to Shh perturbation in the anterior tail of uninjured and injured tadpoles during the stages we are investigating. However, when compared with what we see in the regenerating spinal cord, the domain expansion following treatment with SAG is less extensive, and cyclopamine does not completely ablate expression of Nkx2.2 (Figure 4D). This suggests that while NPC identity remains plastic in the spinal cord of uninjured stage 41 tadpoles and can be modulated by perturbation of Shh, regenerating tissue is the most sensitive to Shh perturbation.

## Discussion

Our principal motivation in undertaking this study was to establish whether the D/V axis of the spinal cord is re-established after injury in *X. tropicalis*, and whether perturbation of Shh signaling influenced that axial patterning. Our study confirms that uninjured, regenerative stage *Xenopus tropicalis* tadpoles maintain expression of developmental markers of D/V NPC domains, and demonstrates that these domains are present during regeneration. We also show that D/V patterning is dependent on Shh signaling in similar manner to embryogenesis, with Shh activating expression of the intermediate and ventral domain transcription factors and inhibiting expression of dorsal markers. In this regard, our results in X. tropicalis align with previous studies done in axolotls^21^. We also demonstrate that regenerating cells show greater sensitivity to Shh perturbations than uninjured cells.

### D/V patterning is dependent on Shh signaling during regeneration

There are at least two mechanisms by which dorsal-ventral patterning information could be propagated from preexisting cells to new cells in regenerating spinal cord. One is that NPCs near the amputation site could acquire a dorsal or ventral identity during development, retain this identity after injury, and undergo mitosis to create progeny with that same positional identity. Alternatively, secreted signals like Shh might be continually required for positional identity in NPCs, and during regeneration these signals would dictate the positional identity of new NPCs, which otherwise would be multipotent. Both mechanisms have been shown to contribute to axolotl regeneration^21,25^.

Our data supports the second model, as we have shown that Shh patterns regenerating NPCs following tail amputation in *Xenopus tropicalis*. Treatment with the Shh inhibitor cyclopamine leads to a significant reduction in Nkx6.1+ cells and the complete absence of Nkx2.2+ cells, which matches with results showing that high and sustained levels of Shh are necessary for the activation of Nkx2.2 expression^39^. Treatment with SAG conversely leads to an increase in Shh activity, causing expansion of Nkx6.1 and Nkx2.2 into more dorsal positions, and the consequent repression of Msx1/2. Unexpectedly, our data showed not only a D/V axis within the regenerating spinal cord, but also an A/P axis of NPC domains. There are a number of signals which may regulate NPC domains orthogonally to the Shh signal. These include posteriorizing signals that are well known to act in spinal cord regeneration, such as FGF and Wnt^40^. Overall, this study demonstrates that, as in the axolotl and in contrast to the anole, *Xenopus* depends upon extrinsic signals to reestablish D/V patterning in the spinal cord after injury. Whether the clonal restriction mechanism utilized in the first model is at all utilized in *Xenopus* remains an intriguing question in the comparative study of regeneration across vertebrate species.

### Species differences in Shh signaling during spinal cord regeneration

The most parsimonious mechanism for the patterning activity we observe would be for Shh to act through Gli-dependent activation of ventral transcription factors and cross-repression of dorsal transcription factors, as thoroughly described in mouse and chick embryogenesis^1,4^. However, recent work in *Xenopus laevis* suggests that Gli1 and Gli2 are dispensable for regenerative growth of the spinal cord^27^. There are several lenses through which we believe our results can be resolved with that study, though these require further testing to fully address. The first possibility, which we consider the most likely, is that the signal transduction mechanism of Shh as a mitogen in spinal cord regeneration may be distinct from its transduction as a morphogen. It may therefore be that Shh acts through Gli-independent non-canonical Ca^2+^ dependent mechanisms to direct growth, while it acts through canonical signaling to establish patterning. We note that the doses of cyclopamine we report here are much lower than those used to repress regenerative growth^27^, and so the sensitivity of each of these pathways to ligand binding may differ. Assessment of spinal cord patterning under Gli-inhibited conditions in *X. laevis* would clarify whether this explanation is most likely. Alternatively, it is plausible that there are species-specific mechanisms in *X. tropicalis* and *X. laevis*, with the latter relying less on canonical Shh signaling than the former. We note that the localization of *shh* transcripts does seem to vary between these species, as elucidated below. Another question for further study is how the perturbation of NPC positional identity by Shh is propagated to post-mitotic neurons and ultimately to spinal cord function. Future work will interrogate the composition and organization of post-mitotic neurons in the regenerating spinal cord, as well as the effects of Shh perturbation on motor function. The source of Shh signal has intriguing differences among regenerative amphibians. During development, both the notochord and the floor plate express *shh* and participate in patterning. However, the axolotl does not regenerate its notochord, leaving the floor plate as the sole source of Shh during regeneration^21^. In contrast, *Xenopus laevis* regenerates the notochord, but it is unclear whether it regenerates the floor plate, as *in situ* hybridization shows no expression of *shh* in the regenerating spinal cord^20,41^, but scRNAseq data shows a *shh*+/*foxa2*+ neural cluster in the regenerate^42^. Here we show that in *Xenopus tropicalis*, a morphologically distinct *shh*-expressing floor plate and notochord are each regenerated, with expression of *shh* detected in each by *in situ* hybridization. As both structures express *shh*, both may contribute as sources of secreted Shh that influence positional identity in the neighboring NPCs of the spinal cord.

### The NPC response to Shh shifts following injury

During development of the non-regenerative mouse and chick, spinal cord maturation is accompanied by a rapid decrease in the number of NPCs lining the central canal. During this time, the Shh signal that was integral to the initial patterning of the neural tube also diminishes, until it finally appears to vanish in the postnatal spinal cord^43,44^. Expression of dorsal-ventral patterning genes is also significantly altered in the mature spinal cord, as expression of Nkx2.2 as well as dorsal markers diminish and the entire ependyma becomes Nkx6.1+^44,45^. In contrast to these observations in non-regenerative amniotes, adult axolotls retain expression of Shh in the floor plate ^21,24^. We find that *Xenopus tropicalis* tadpoles, like axolotls, maintain embryonic patterning factors in the spinal cord in regenerative-stage tadpoles. The maintenance of progenitor domains may be related to the fact that anamniotes like *Xenopus*, axolotls, and zebrafish must undergo a second round of neurogenesis during their larval stages to generate adult neuronal types^46^.

We demonstrate that Shh continues to be expressed in the notochord and floor plate up until tadpole stages, and that Nkx2.2+, Nkx6.1+, and Msx1/2+ domains retain their embryonic positions through these stages as well. We also present evidence that the anterior tissue in both uninjured and injured tadpoles remains sensitive to Shh activation or inhibition, with greater sensitivity in regenerating NPCs. Sensitivity to Shh perturbation subsequently declines until 3 dpa, after which there is no change to the positioning of the Nkx6.1 domain in response to cyclopamine or SAG treatment. This could represent the progenitors reaching a more mature state and corresponds roughly with the onset of neurogenesis in the anterior regenerate. Both the ongoing sensitivity of NPC domains to Shh perturbation and the continued expression of Shh past its initial role in dorsal-ventral patterning have interesting implications for the role in cell fate plasticity of the *X. tropicalis* spinal cord. *Xenopus* are notable in that regenerative competence declines with age, and so one aspect of regenerative competence may be tied to the ability of regenerative NPCs to respond to Shh. Future work will interrogate the plasticity of NPC fate in non-regenerative stages.

## Methods

### *Xenopus tropicalis* amputation assay

NF stage 41 tadpoles were anesthetized with 0.05% MS-222 in 1/9x MR and tested for response to touch prior to amputation surgery. Once fully anesthetized, a sterilized scalpel was used to amputate the posterior third of the tail. Amputated tadpoles were removed from anesthetic media within 10 min of amputation into new 1/9x MR. Tadpoles were kept at a density of no more than 2.5 tadpoles per mL. Tadpoles were fixed for 1 h in 1x MEM with 3.7% formaldehyde at room temperature at 2 dpa (days post amputation), 3 dpa, or 4 dpa.

#### Immunohistochemistry

For whole mount immunohistochemistry, fixed tadpoles were permeabilized by washing 3 × 15 min in PBS +0.01% Triton X-100 (PBT) with the exception of tadpoles stained for Nkx2.2, which were washed with 0.05% Triton X-100 for the entire protocol. Tadpoles were blocked for 1 h at room temperature in 10% CAS-block (Invitrogen #00-8120) in PBT, with the exception of tadpoles stained for Msx1/2 which were incubated at 4°C overnight in 10% CAS-block. Then, tadpoles were incubated in primary antibody [1:10 mouse Msx1/2 (4G1, DSHB); 1:10 mouse Nkx2.2 (74.5A5, DSHB); 1:50 mouse Nkx6.1 (F55A10 , DSHB)] diluted in 100% CAS-block overnight at 4°C. Tadpoles were then washed 3 × 10 min at room temperature in PBT and blocked for 30 min in 10% CAS-block in PBT. Secondary antibody (goat anti-mouse 594, ThermoFisher A11032) was diluted 1:500 in 100% CAS-block and incubated for 2 h at room temperature. Tadpoles were then washed 3 × 10 min in PBT followed by a 10-min incubation in 1:2000 DAPI (Sigma D9542) before being washed with 1xPBS for 10 more minutes. Isolated tails were mounted on slides in ProLong Diamond (ThermoFisher P36961). For cryosectioned immunohistochemistry, fixed tadpoles were washed with 3 x 15 min 1X PBS, 1 x 15 min 15% sucrose in 1X PBS, then overnight at 4°C. Tadpoles were then transferred into OCT (Thermofisher 23-750-571) and kept at -80°C until sectioning with a Leica CM3050 S Cryostat. Immunohistochemistry proceeded as described above. Images were acquired using a Leica DM 5500 B microscope using a 20X or 40X objective and processed using FIJI image analysis software.

### Whole mount in situ hybridization

Embryos and tadpoles were fixed overnight in 1x MEM with 3.7% formaldehyde at 4°C. Xenopus tropicalis multibasket in situ hybridization protocols were followed as described^47^, with the notable change that pre-hybridization was always performed overnight. Isolated tails were mounted on slides in ProLong Diamond (ThermoFisher P36961) and imaged on a Leica M205 FA with a color camera. For cross-sections, tadpoles were stained then cryosectioned as described above. Probes were synthesized using the following primer pairs designed against a single exon of each mRNA: shh (forward – gctttctacacctgccttgc, reverse – taatacgactcactatagggtgagtcatgagtcggtctgc), foxa2 (forward – aactgcagcagctttggaa, reverse – taatacgactcactatagggcccgtttaagagtaagaactgagg. RefSeq annotation derived from Xenbase (http://www.xenbase.org/, RRID:SCR_003280)^48^ genome v10.0.

#### Pharmacological Inhibition

Cyclopamine (Cayman Chemical, 11321) was resuspended to a 10 mM stock in DMSO, and SAG (GLPBio, GC50135) to a 5 mM stock in DMSO. Uninjured and injured tadpoles were reared with 0.01% DMSO, 500 nM SAG or 1 μM cyclopamine diluted in 1/9x MR until collection at 2 dpa, 3 dpa, or 4 dpa following treatment. Other concentrations were used in establishing doses and are reported in Figure S2.

### Quantification and statistical analysis

Whole mount immunostained tail images were processed in Fiji^49^ and the regenerate spinal cord was isolated using the Straighten tool. Segmentation, counting and measurements of Nkx6.1+/Nkx2.2+/Msx+ cells was done using the Fiji Analyze Particles tool. Dorsal/Ventral position after DMSO, cyclopamine, or SAG treatment were compared using one-way ANOVA followed by Tukey’s multiple comparisons test or pairwise Wilcoxon two-sample tests depending on normality of data. The Shapiro-Wilk test was used to assess normality. N (number of animals) has been reported for regeneration experiments on graphs. The data for each marker consists of at least two independent experiments. Information regarding statistics can be found in the figure legend or on the figure panel with the data. Boxplots and stacked bar plots were generated using the R package ggplot2^50^.

## Acknowledgements

We thank members of the Wills lab for critical comments during the preparation of this manuscript and support in frog husbandry. We thank Laura Borodinsky and Andrew Hamilton for feedback on a preliminary paper draft. We thank the Reh and Kong lab for access and training on their cryostats. We thank Jennifer Kong for consultation about the use of SAG. We thank Xenbase for curation of genomic and literature information and the National Xenopus Resource for frogs. AAS was supported by the Cellular and Molecular Biology Training Grant NRSA 5T32GM136534-02 from NIGMS. SBP was supported by a Mary Gates Fellowship for Undergraduate Research. This work was supported by NINDS R01NS099124 to AEW.

---

**Figure S1.**
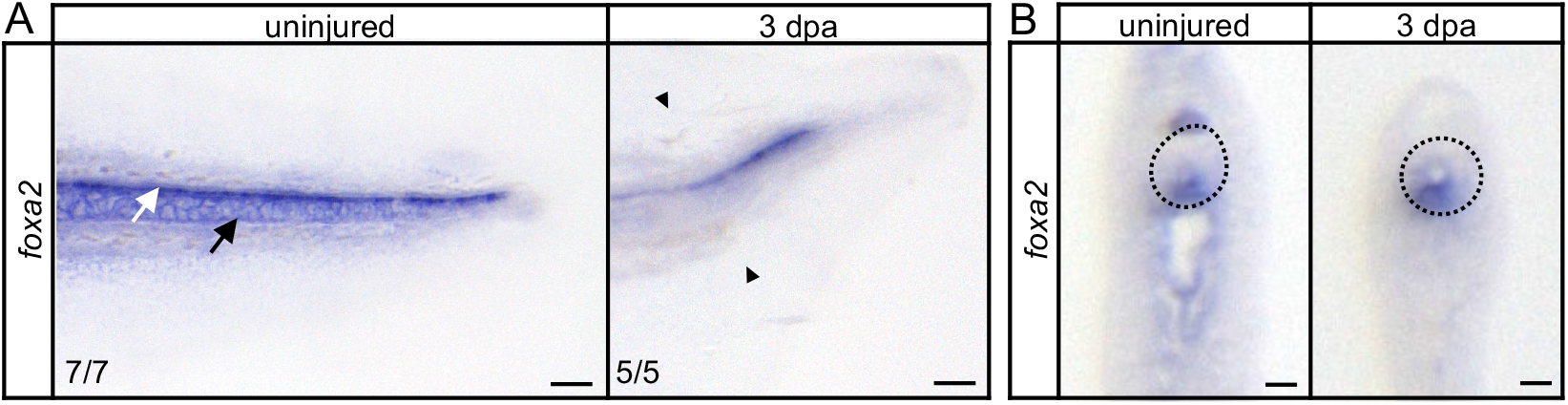
foxa2 is present within the regenerating spinal cord floor plate. Images of *in situ* hybridization visualizing *foxa2* expression at uninjured and 3 dpa in lateral view (**A**) and transverse (**B**). **A**. White arrow indicates spinal cord floor plate, black arrow indicates notochord. Scale bar = 100 µm. Paired triangles represent amputation plane. n/n is noted. **B**. Dotted line represents spinal cord border. Scale bar = 10 µm.

**Figure S2.**
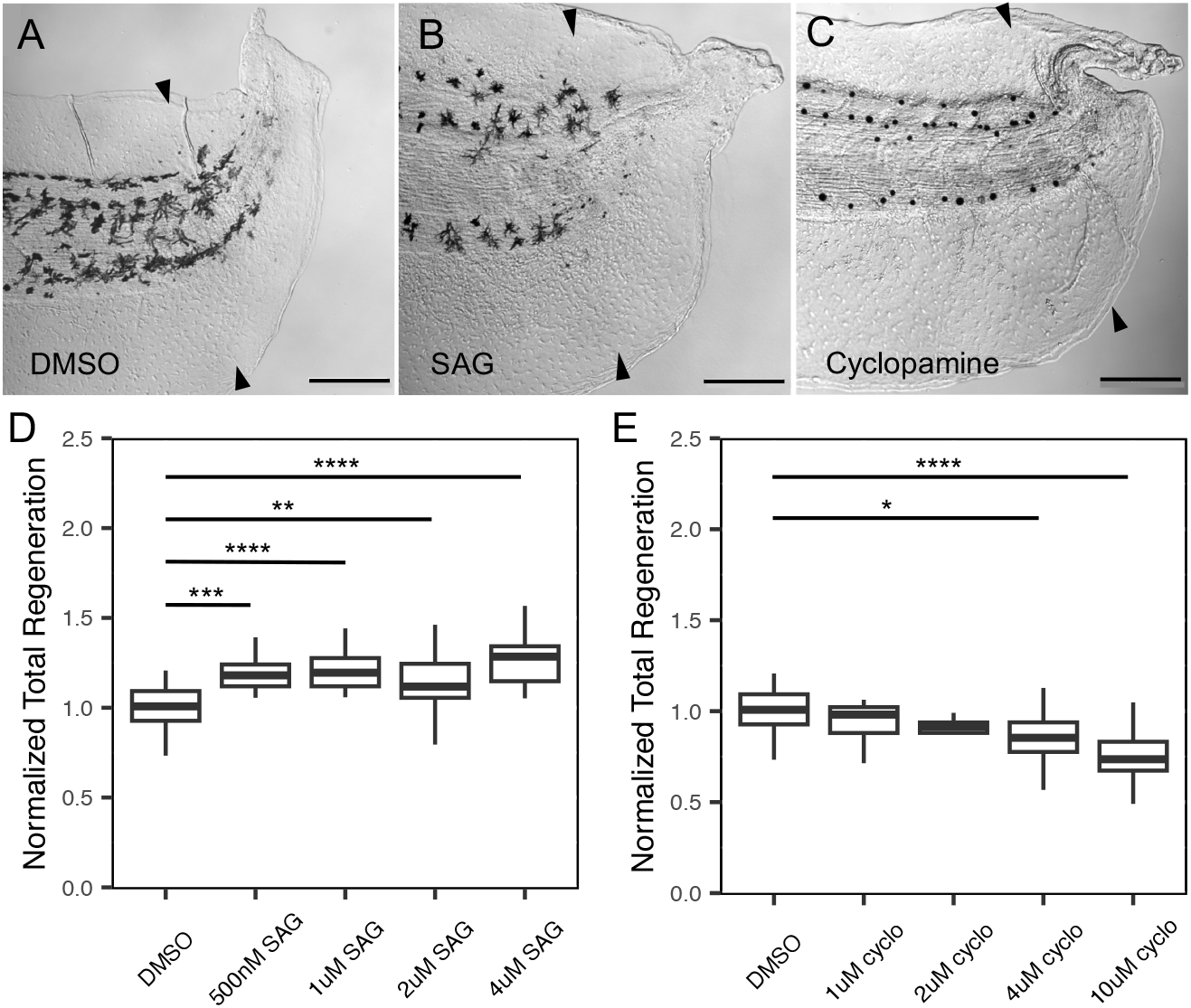
Shh perturbation alters total regeneration. (**A-C**) DIC images of 3 dpa regenerated tails following treatment with (**A**) DMSO (**B**) 4 µM SAG (**C**) 10 µM Cyclopamine. Paired triangles represent amputation plane. Scale bar = 200 µm. (**D-E**) Dose curves with total regeneration at 3 dpa normalized to DMSO treated group within respective clutch for (**D**) SAG and (**E**) cyclopamine. Significance was determined by one-way ANOVA followed by Tukey’s multiple comparisons test. (^*^ p<0.05, ^**^ p<0.01, ^***^ p<0.001, ^****^ p<0.0001).

**Figure S3.**
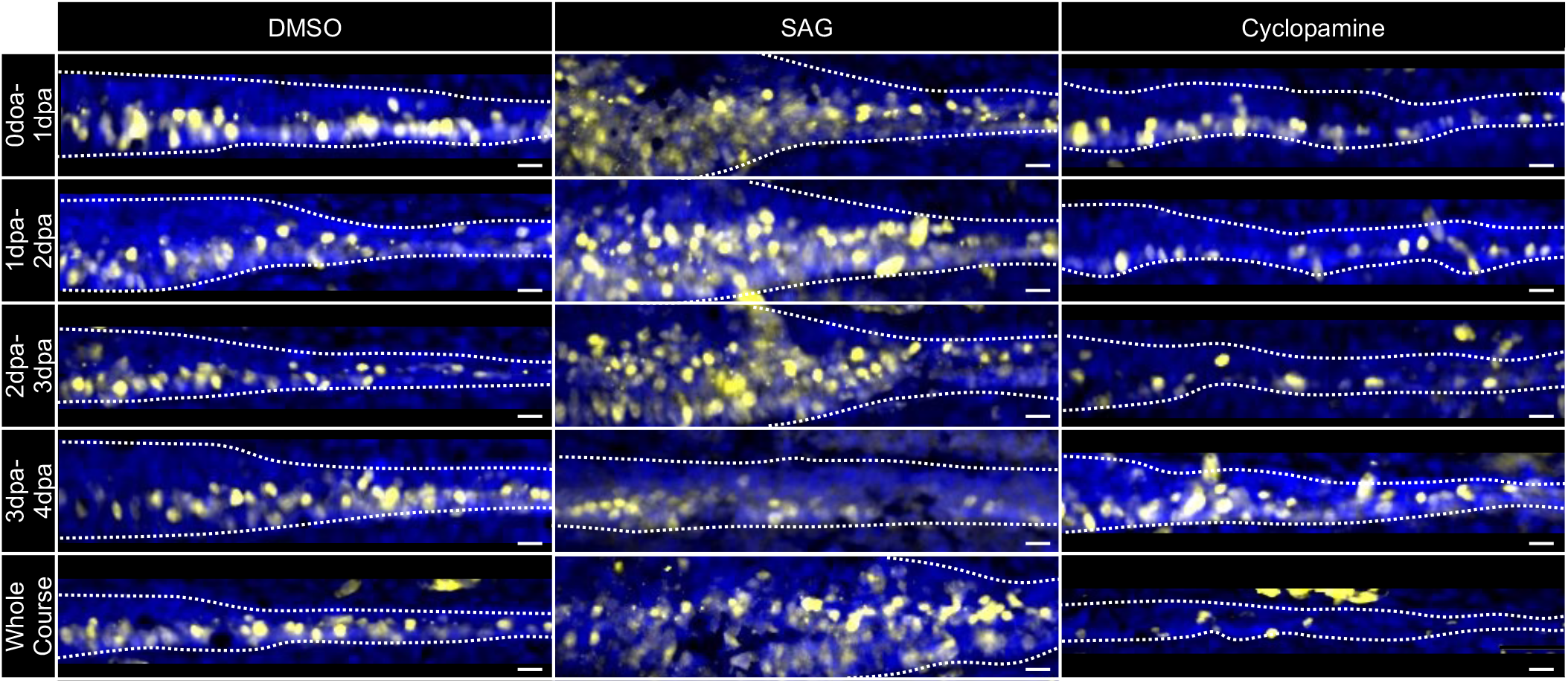
Regenerating NPCs are more sensitive to Shh modulation before 3 dpa. Images of Nkx6.1+ cells after treatment with DMSO, SAG, or cyclopamine during specific regenerative timepoints. Dotted white lines represent spinal cord boundary. Scale bar = 10 µm.

